# Calorie Restriction Outperforms Bariatric Surgery in a Murine Model of Obesity and Triple-Negative Breast Cancer

**DOI:** 10.1101/2023.05.19.541484

**Authors:** Michael F Coleman, Kristina K Camp, Tori L McFarlane, Steven S Doerstling, Subreen A Khatib, Erika T Rezeli, Alfor G Lewis, Alex J Pfeil, Laura A Smith, Laura W Bowers, Farnaz Fouladi, Weida Gong, Elaine M Glenny, Joel S Parker, Ginger L Milne, Ian M Carroll, Anthony A Fodor, Randy J Seeley, Stephen D Hursting

## Abstract

Obesity promotes triple-negative breast cancer (TNBC), and effective interventions are urgently needed to break the obesity-TNBC link. Epidemiologic studies indicate that bariatric surgery reduces TNBC risk, while evidence is limited or conflicted for weight loss via low-fat diet (LFD) or calorie restriction (CR). Using a murine model of obesity-driven TNBC, we compared the antitumor effects of vertical sleeve gastrectomy (VSG) with LFD, chronic CR, and intermittent CR. Each intervention generated weight and fat loss and suppressed tumor growth relative to obese mice (greatest suppression with CR). VSG and CR regimens exerted both similar and unique effects, as assessed using multi-omic approaches, in reversing obesity-associated transcriptional, epigenetic, secretome, and microbiota changes and restoring antitumor immunity. Thus, in a murine model of TNBC, bariatric surgery and CR each reverse obesity-driven tumor growth via shared and distinct antitumor mechanisms, and CR is superior to VSG in reversing obesity’s procancer effects.

## Introduction

Obesity is an established risk factor for the development and progression of several breast cancer subtypes (1). This includes triple-negative breast cancer (TNBC), an aggressive and particularly deadly breast cancer subtype (2). The prevalence of obesity in American women now exceeds 40% (3), and effective interventions to lessen the procancer effects of obesity are urgently needed (4–6). Unfortunately, the current literature regarding the reversibility of the obesity-breast cancer link via weight loss interventions is limited. Epidemiologic studies clearly show that bariatric surgery reduces the risk of numerous obesity-associated cancers, including breast cancer (7–10). The most common weight loss surgery in the US and worldwide is vertical sleeve gastrectomy (VSG) (11, 12). However, all types of bariatric surgery carry surgical complication risks, are very expensive, have strict exclusion criteria, and hence are unavailable to most women with obesity (13). Dietary interventions, which in theory represent widely available and inexpensive means through which weight loss may be achieved, are difficult for many people with obesity to sustain (14–17). Moreover, the epidemiologic literature is limited and inconsistent regarding the ability of dietary weight loss to reverse breast cancer risk or progression in women with obesity (18).

Obesity promotes breast cancer progression through multiple interacting mechanisms, including chronic inflammation. Indeed, obesity alters gene expression and dysregulates systemic adipokines, cytokines, prostaglandins, oxylipins, and other inflammatory markers to promote immunosuppression and procancer signaling in the tumor microenvironment (2, 19). Obesity-associated changes in gut microbial communities appear to contribute to the inflammatory and immune alterations via microbe-derived metabolites (20). Successful, sustained weight loss interventions can modulate many of these mechanisms (5, 21–24), although their causal relationships with reduced breast cancer burden remain unclear (25) (26).

Low-fat diet (LFD) regimens are the most commonly studied weight loss interventions in people with obesity, but exhibit consistently poor long-term weight loss success (27).

Calorie restriction (CR) is widely effective at reducing obesity-driven tumor incidence and progression in preclinical studies (1, 5) but difficult to implement in people.

Intermittent CR (ICR) regimens, where periods of low-calorie consumption are interspersed with unrestricted eating, have emerged as effective weight loss interventions and are more easily implemented than chronic CR (CCR) (28).

Studies in obese animals that characterize the antitumor effects and mechanisms of weight loss achieved via bariatric surgery as compared with calorie restriction interventions (CCR or ICR regimens) have not yet been reported. We partially addressed this gap in a recent report—using the same murine TNBC model and dietary weight loss interventions studied here—and established that the genomic, epigenetic, and antitumor effects of obesity are reversible by CCR and ICR regimens, and, to a lesser extent, LFD (5). Herein, we confirm and extend those findings to identify the shared and distinct antitumor effects and mechanisms of VSG relative to LFD, CCR, or ICR in a mouse model of obesity and TNBC. Specifically, we conducted two preclinical studies to test the hypotheses that i) weight loss induced by VSG or CR interventions reverses obesity-driven TNBC progression by remodeling the transcriptomic, epigenetic and immune landscape of tumors and adjacent adipose; and ii) weight loss-mediated suppression of tumor growth is concomitant with alteration of intestinal microbiomes, circulating cytokines, and oxylipins.

## Results

### Bariatric surgery and LFD each promote weight loss in obese mice, limit obesity-driven TNBC growth, and promote markers of antitumor immunity

In Study 1 (**Fig. 1A**) we determined whether obesity-driven mammary cancer could be limited by weight loss resulting from VSG or LFD dietary change. Prior to surgery, DIO mice weighed significantly more than CON mice (**Fig. 1B**). While all DIO mice exhibited some weight loss immediately after surgery, only DIO-VSG and DIO-LFD groups sustained this weight loss (**Fig. 1C, Fig. S1A**). Prior to tumor cell injection and at the end of the study, DIO-HFD mice remained heavier than all other groups and DIO-LFD mice weighed more than CON-LFD mice (**Fig. 1D-E**). Similarly, ex vivo tumor mass was greater in the DIO-HFD group relative to all other groups (**Fig. 1F**). Tumor mass from DIO-VSG mice was not different to that of CON-LFD or DIO-LFD mice (**Fig. 1F**).

**Figure 1.**
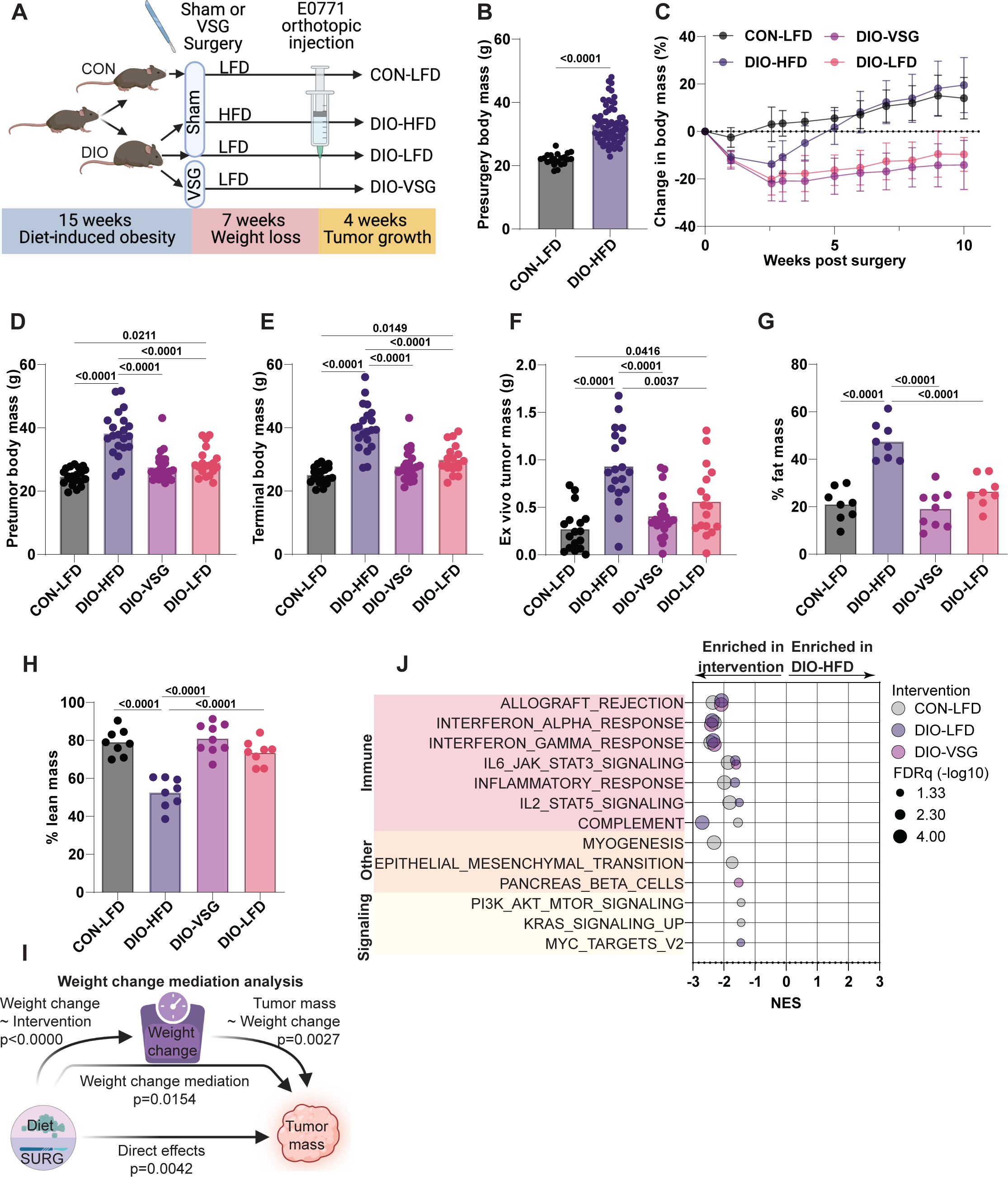
Dietary and surgical weight loss blunt obesity-driven tumor growth (A) Schematic of study design. (B) Body mass prior to weight loss interventions. (C) Change in body weight over time following weight-loss interventions. (D) Body mass prior to tumor cell injection. (E) Terminal body mass. (F) Ex vivo tumor mass. (G-H) Body composition following weight loss interventions. (I) Mediation analysis of weight change following weight-loss intervention on tumor mass. (J) Hallmark gene sets determined significant by GSEA of tumor transcriptomics in pairwise comparisons with DIO-HFD. Gene sets grouped and colored as immune, other, and signaling-related. (B- F, I) n=21 CON-LFD, 21 DIO-HFD, 24 DIO-VSG, 19 DIO-LFD. (G-H) n=8 CON-LFD, 8 DIO-HFD, 9 DIO-VSG, 8 DIO-LFD. (J) n=6 CON-LFD, 6 DIO-HFD, 6 DIO-VSG, 5 DIO-LFD. (B-H) One-way ANOVA with Tukey’s post hoc test.

Tumors mass was greater in DIO-LFD mice relative to CON-LFD mice (**Fig. 1F**).

We next sought to understand how body composition was altered by VSG and LFD interventions. MRI showed that, relative to the DIO-HFD regimen, each weight loss intervention resulted in reduced relative body fat and increased relative lean mass (**Fig. 1G-H**). Paralleling this remodeling of adipose tissue, we found that leptin, resistin, GLP1, glucagon, TNF⍺, and IL6 were elevated in serum from DIO-HFD mice relative to CON-LFD, and DIO-VSG suppressed this obesity-driven increase (**Fig. S2A-F**). Serum from DIO-LFD mice had a similar overall suppression of obesity-driven elevation of adipokines except glucagon and TNFɑ (**Fig. S2A-F**).

Given the similar profile of terminal body weight and tumor mass (**Fig. 1E-F**), we used mediation analysis to determine if changes in tumor mass could be explained by proportion of weight lost during the 11 weeks between pre-intervention and study end point. We employed mediation analysis to quantify the potential for intervention-driven weight loss to explain terminal tumor mass. Tumor mass was, in part, explained by the percent of weight lost between weight loss interventions and study end (p=0.0154) and in part by weight change-independent sources of variance (direct effects) (p = 0.004) (**Fig. 1I**).

To better understand how these weight loss interventions altered tumor growth, we performed transcriptomic profiling of tumors using Affymetrix microarray, followed by GSEA (29) using MSigDB Hallmark gene sets (30). Relative to tumors from CON-LFD mice, tumors from DIO-HFD mice exhibited suppression of several immune, differentiation, and signaling-related gene sets (**Fig. 1J**). Obesity-driven suppression of immune-related gene sets was effectively reversed by both DIO-VSG and DIO-LFD weight loss interventions (**Fig. 1J**). To gain additional insight into the processes altered in tumor transcriptomic profiles, we performed GSEA using Gene Ontology Biological Processes (GOBP) (31) gene sets. To limit redundancy, we subjected all significantly enriched gene sets to enrichment mapping (32) and clustered by similarity coefficient to identify commonly enriched processes and themes. Immune-related signaling dominated the clusters identified for each binary comparison of DIO-HFD with CON- LFD, DIO-VSG, and DIO-LFD (**Fig. S3A-C**).

### Bariatric surgery and dietary weight loss drive distinct transcriptomic changes in mammary adipose tissue

To understand molecular alterations in adipose tissue gene expression accompanying the changes in body composition, we performed RNAseq transcriptomic profiling on mammary adipose tissue contralateral to the tumor. The comparisons of DIO-HFD versus CON-LFD, DIO-HFD versus DIO-VSG, and DIO-HFD versus DIO-LFD revealed numerous differentially expressed genes characterized by a preponderance of overexpressed genes in DIO-VSG relative to DIO-HFD, and suppression of gene expression in CON-LFD and DIO-LFD relative to DIO-HFD (**Fig. 2A-D**).

**Figure 2.**
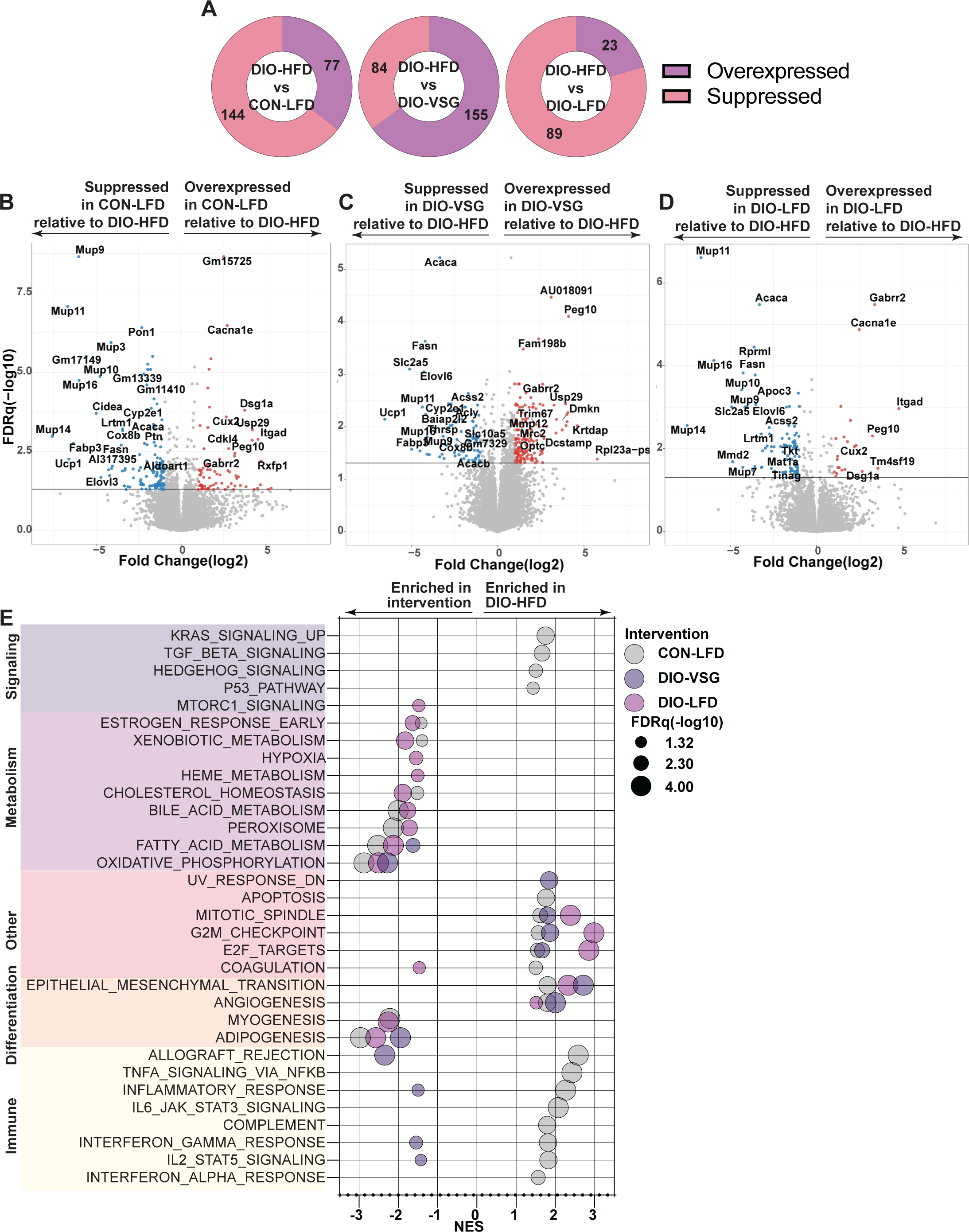
Transcriptomic analysis of mammary adipose tissue following dietary and surgical weight loss reveals discordant metabolic and immune signaling (A) Distribution of DEG relative to DIO-HFD. (B-D) Volcano plots of DEG generated in the comparison between CON-LFD, DIO-VSG, and DIO-LFD relative to DIO-HFD, respectively. (E) Hallmark gene sets determined significant by GSEA of adipose tissue transcriptomics in pairwise comparisons with DIO-HFD. Gene sets grouped and colored as signaling, metabolism, other, differentiation, and immune-related. n=4/group.

In contrast to the tumor transcriptomic profiles, GSEA of RNAseq data revealed marked enrichment of inflammatory signaling in the mammary tissue from DIO-HFD mice relative to CON-LFD (**Fig. 2E**). Relative to DIO-HFD, weight loss by diet did not alter inflammatory signaling, and VSG only modestly altered inflammatory signaling (**Fig. 2E**). Numerous obesity-driven changes in markers of growth/survival, metabolism, and differentiation were found in the comparison of adipose tissue from DIO-HFD mice with that of CON-LFD mice (**Fig. 2E**). Adipose tissue from DIO-VSG and DIO-LFD mice showed restoration of many of these obesity-driven pathway alterations when compared with adipose tissue from DIO-HFD mice (**Fig. 2E**).

### Bariatric surgery, but not LFD-driven weight loss, drives epigenetic alterations concordant with predicted mediators of transcriptomic changes in adipose from humans who underwent bariatric surgery

We hypothesized that obesity and weight loss would promote transcriptomic reprogramming in part via changes in epigenetic regulation through DNA methylation. To test this hypothesis, we performed reduced-representation bisulfite sequencing on DNA isolated from mammary adipose tissue contralateral to tumor. Relative to DIO-HFD adipose tissue, CON-LFD adipose tissue contained 839 differentially methylated genes (DMG), DIO-LFD adipose tissue contained 1,062 DMG, and DIO-VSG adipose tissue contained 31,424 DMG (**Fig. 3A-D**). To identify transcription factors predicted to regulate these DMG, GSEA was performed on each pairwise comparison. Only the comparison of DIO-HFD and DIO-VSG demonstrated significant enrichment of transcription factors including NFKB1, BACH1, and FXR1 (**Fig. 3E**). Finally, we sought to determine if the transcription factors that were responsive to DIO-VSG-driven epigenetic changes were also responsive in human adipose tissue following bariatric surgery. We performed RegEnrich analysis of GSE59034 to identify likely transcriptional mediators of adipose remodeling in humans following bariatric surgery and found all three transcription factors (NFKB1, BACH1, FXR1) were significantly associated with gene expression changes following bariatric surgery (**Fig. 3F**).

**Figure 3.**
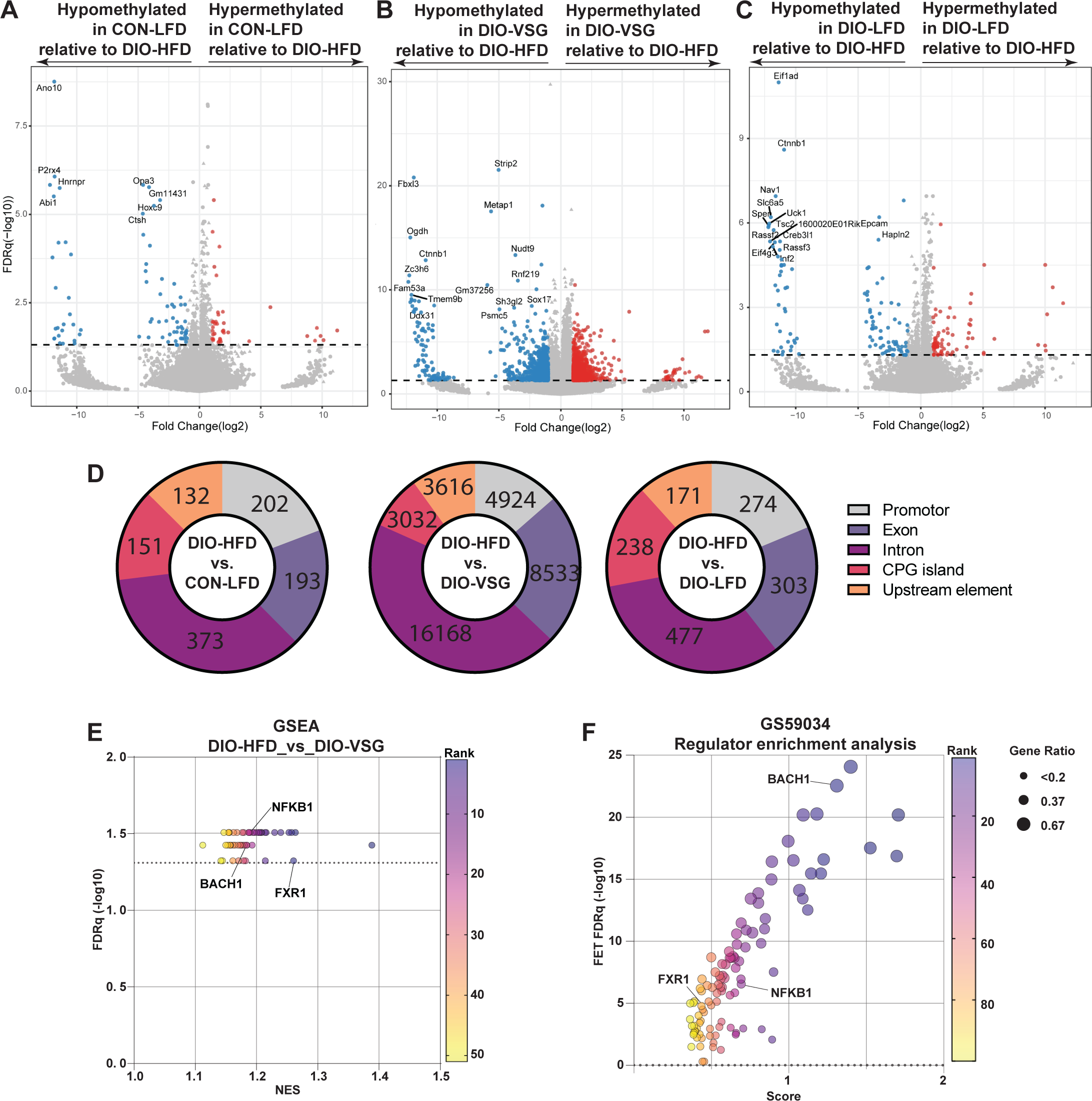
Epigenetic regulation through DNA methylation of mammary adipose tissue reveals transcriptional mediators of the gene expression profile conserved between human and mouse adipose tissue following surgical but not dietary weight loss (A-C) Volcano plots of DMG generated in the comparison between CON-LFD, DIO- VSG, and DIO-LFD relative to DIO-HFD, respectively. (D) Distribution of DMG relative to DIO-HFD. (E) MSigDB C3 gene sets determined significant by methylGSA of adipose tissue RRBS in comparison of DIO-VSG with DIO-HFD. (F) Regulator enrichment analysis of adipose tissue from patients who were never obese or obese pre/post bariatric surgery (GSE59034). (A-E) n=4/group, (F) n=16/group.

### Remodeling of fecal microbiotas is associated with weight loss and tumor size

Using amplicon sequencing of 16S rDNA isolated from fecal matter, we first confirmed that CON-LFD and DIO-HFD microbiotas were distinct. DIO-HFD microbial communities had fewer observed sequence variants relative to the CON-LFD group while Shannon index was unchanged (**Fig. S4A-B**). β-diversity was also distinct between CON-LFD and DIO-HFD microbiotas as illustrated by changes in proportional abundances of the ten most prevalent genera and significant separation of groups by nonmetric multidimensional scaling (NMDS) plot of Bray-Curtis distances (**Fig. S4C-D**). Both weight loss interventions and DIO-HFD reduced the number of observed sequence variants but not Shannon index (which accounts for evenness of microbial composition) relative to CON-LFD associated microbial communities (**Fig. S4E-F**). The proportional abundances of genera in fecal samples from both weight loss groups were more similar to one another than to other groups (**Fig. S4G**). NMDS plot of Bray-Curtis distances revealed close clustering of microbial communities among the weight loss interventions, of which both DIO-VSG and DIO-LFD associated communities were different to the fecal microbiota in CON-LFD mice. Further, fecal microbiotas from DIO-LFD were distinct to those from DIO-HFD mice (**Fig. S4H**). Finally, we tested the association of each genus with percent body weight change and terminal tumor mass. While several genera were significantly associated with these percent body weight change, no genera were significantly associated with tumor mass (**Fig. S4I-K**).

### Enhanced weight loss through either CCR or ICR more effectively limits obesity-driven TNBC growth than bariatric surgery and restores markers of antitumor immunity

In Study 1, weight loss was a significant mediator of smaller tumor size in both the VSG- and LFD-induced weight loss groups and potently remodeled adipose tissue in DIO- VSG mice. Hence, we sought in Study 2 (**Fig. 4A**) to determine if greater weight loss achieved through CCR or ICR would mimic or surpass bariatric surgery in limiting mammary cancer growth. As expected, DIO mice weighed significantly more than CON mice prior to weight loss (**Fig. 4B**). While all weight loss interventions promoted significant weight loss, both DIO-CCR and DIO-ICR promoted greater and more sustained weight loss than DIO-VSG (**Fig. 4C-E**).

**Figure 4.**
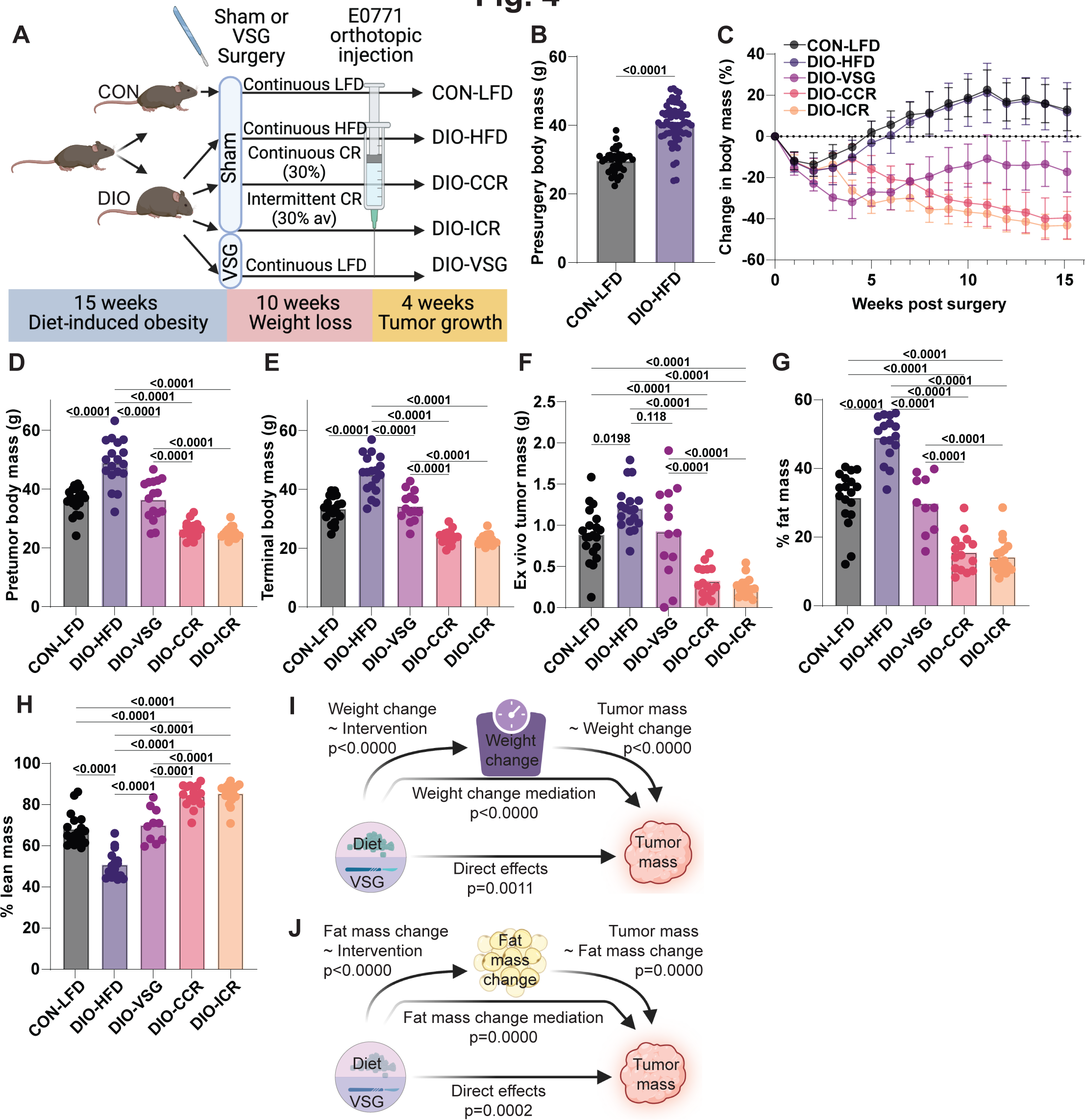
Dietary weight loss via caloric restriction outperforms surgical weight loss to blunt obesity-driven tumor growth (A) Schematic of study design. (B) Body mass prior to weight loss interventions. (C) Change in body weight over time following weight-loss interventions. (D) Body mass prior to tumor cell injection. (E) Terminal body mass. (F) Ex vivo tumor mass. (G-H) Body composition following weight loss interventions. (I) Mediation analysis of weight change following weight-loss intervention on tumor mass. (J) Mediation analysis of fat mass change following weight-loss intervention on tumor mass. n=20 CON-LFD, 18 DIO-HFD, 14 DIO-VSG, 19 DIO-ICR, 16 DIO-CCR. (B-H) One-way ANOVA with Tukey’s post hoc test.

Relative to CON-LFD mice, DIO-HFD mice had significantly larger tumors (**Fig. 4F**). The mean tumor mass of DIO-VSG mice (0.92 ± 0.56 g) was not significantly different than either CON-LFD (0.88 ± 0.32 g) or DIO-HFD (1.20 ± 0.27 g) (**Fig. 4F**). In contrast, mice from both calorie restriction interventions—DIO-CCR and DIO-ICR—had significantly smaller tumors relative to all other groups (**Fig. 4F**). At study termination, DIO-HFD mice had a higher proportion of body mass accounted for by fat mass, and a lower proportion by lean mass, than mice from all other groups (**Fig. 4G-H**). Bariatric surgery changed the composition of lean and fat mass to be not different to CON-LFD mice (**Fig. 4G-H**). Chronic and intermittent calorie restriction both further reduced percent fat mass and increased percent lean mass relative to all other groups (**Fig. 4G-H**). Similar to Study 1, we identified body weight loss (p<0.0001), in addition to weight loss-independent effects (p<0.01), as significant mediators of the antitumor effects resulting from the weight loss interventions in Study 2 (**Fig.4I**). We identified changes in fat mass (p<.001) and fat mass loss-independent effects (p<0.05) as significant mediators of the antitumor effects of the tested interventions (**Fig. 4J**).

To understand signaling and pathways altered by the weight loss interventions, we performed tumor transcriptomic profiling followed by GSEA. Concordant with findings from Study 1, we identified marked suppression of immune related gene sets in tumors from DIO-HFD mice relative to CON-LFD (**Fig. 5A**). All weight loss interventions (DIO- VSG, DIO-CCR, DIO-ICR) reversed tumoral immunosuppression, as indicated by enrichment of immune related gene sets relative to DIO-HFD (**Fig. 5A**).

**Figure 5.**
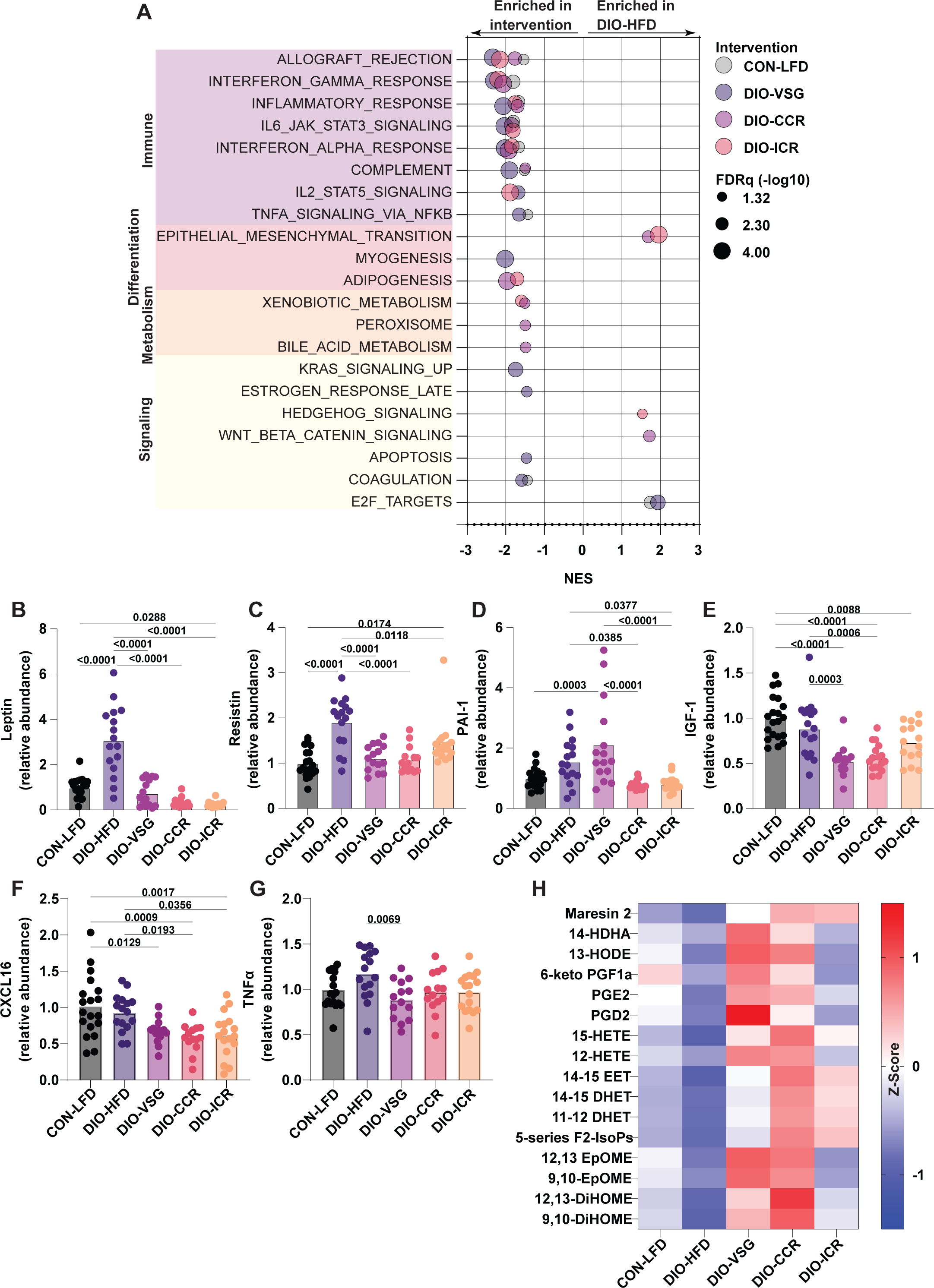
Body weight loss and adiposity mediate blunting of obesity-driven tumor growth by dietary and surgical weight loss interventions (A) Significant GSEA Hallmark gene sets for tumor transcriptomic data in pairwise comparisons with DIO-HFD. (B-G) Circulating adipokines determined by multiplex ELISA. (H) Mammary adipose tissue oxylipin levels determined by UPLC-MS (Z-Score). (A) n=6/group. (B-G) n=19 CON-LFD, 16 DIO-HFD, 14 DIO-VSG, 14 DIO-ICR, 17 DIO-CCR. (H) n=11 CON-LFD, 10 DIO-HFD, 9 DIO-VSG, 10 DIO-ICR, 13 DIO-CCR.

Given the prominent remodeling of obesity-driven adipose tissue dysfunction demonstrated in Study 1, we next confirmed whether similar alterations of circulating adipokines was achieved following either dietary or surgical weight loss in Study 2. Serum leptin and resistin levels were increased in DIO-HFD relative to CON-LFD mice (**Fig. 5B-C**). Both surgical and dietary weight loss reverted obesity-driven elevation of leptin and resistin levels, while only dietary weight loss reverted elevation of PAI-1 (**Fig. 5B-D**). Serum IGF-1 was reduced in DIO-VSG, DIO-CCR, and DIO-ICR mice relative to CON-LFD mice (**Fig. 5E**). Serum CXCL16 was reduced in all weight loss intervention groups relative to the CON-LFD group (**Fig. 5F**). Serum TNF⍺ was reduced in mice that underwent VSG relative to DIO-HFD mice (**Fig. 5G)**. We also found overall suppression of oxylipins in adipose tissue from DIO-HFD mice relative to other groups, particularly DIO-VSG and DIO-CCR (**Fig. 5H)**.

### Sustained caloric restriction and bariatric surgery promote distinct changes in cecal microbial communities

To assess whether diet-induced obesity or weight loss in Study 2 altered cecal microbial communities, we performed 16S rDNA amplicon sequencing. We found overall limited changes in ⍺-diversity measures of microbial communities across groups as determined by the number of observed sequence variants and Shannon index (**Fig. 6A-B**). β- diversity of cecal microbiotas was significantly different between all intervention groups, illustrated by NMDS plot of Bray-Curtis distances and the taxonomic profiles illustrating the top ten identified genera (**Fig. 6C-D**). Additionally, we used Spearman correlation to identify genera significantly associated with percent body weight change and tumor mass (**Fig. 6E-G**). Among numerous significant associations, only *Hungatella* was significantly associated with both tumor mass and percent body weight loss.

**Figure 6.**
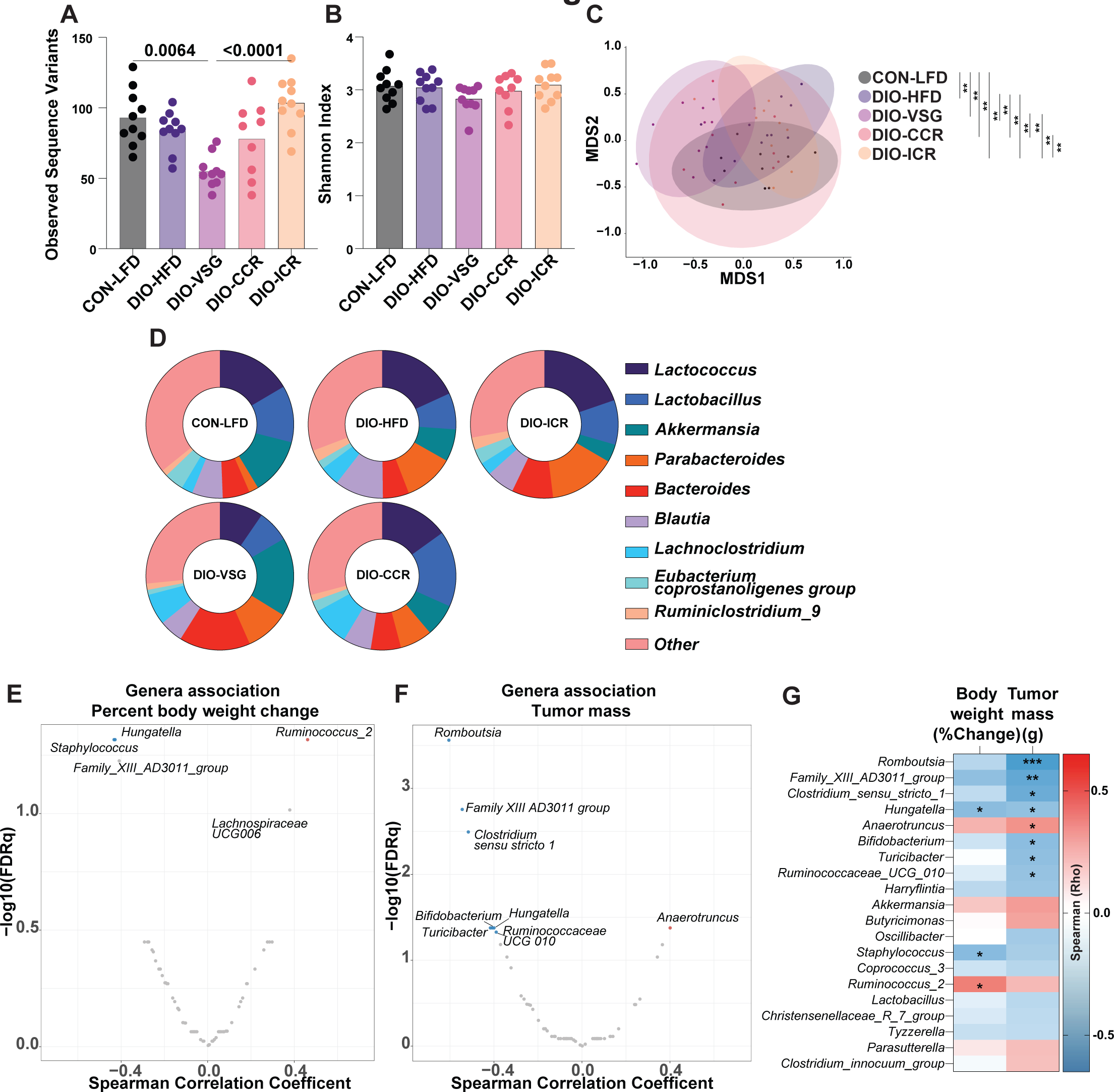
Cecal *Hungatella* abundance associates with both body weight loss and tumor mass (A) Observed sequence variants. (B) Shannon index. (C) NMDS plot of Bray-Curtis distances. (D) Relative contribution of the 10 most frequent genera to each group. Spearman correlation between all genera and (E) percent body weight change and (F) tumor mass. (G) Spearman correlation coefficients of the 20 genera showing the highest correlation coefficients with percent body weight change and tumor mass. n=10/group.

## Discussion

Herein we demonstrate that VSG and CR reverse obesity-driven tumor growth via both distinct and shared mechanisms in a murine model of TNBC. Obesity-associated changes in transcriptional, epigenetic, secretome, intestinal microbiota and antitumor immunity all exhibited both redundant and distinct responses to VSG and CR.

We and others have established that obesity-driven immune cell dysfunction drives immunosuppression in the tumor microenvironment that can be reverted by CR regimens (5, 21–24). Further, recent work has demonstrated an association between bariatric surgery and restoration of antitumor immunity (33). We identified shared and unique transcriptional signatures indicative of restoration of antitumor immunity in mice receiving the VSG and CR regimens. These data are congruent with a sizable literature supporting an immunomodulatory effect of bariatric surgery in both preclinical and patient-derived samples, principally in adipose tissue and not in the context of breast cancer (4). Indeed, we demonstrate extensive adipose reprogramming following weight loss, and identify FXR1, NFKB1, and BACH1 as candidate transcriptional regulators that are common to both human and mouse adipose tissue reprogramming following surgical, but not dietary, weight loss. FXR1, NFKB1, and BACH1 are potent regulators of obesity-driven inflammation and metabolic dysfunction (34–36).

Using mediation analyses we also found that the extent of total body weight loss or adipose weight loss, regardless of intervention, significantly mediates the reduction in tumor growth. This conclusion is supported by substantial literature indicating that obesity-driven adipose inflammation promotes the protumor effects of obesity in models of breast cancer (37).

We identified the cecal microbial genera *Hungatella* to be associated with both reduced tumor growth and body weight loss. Previous reports have shown that both dietary and surgical weight loss interventions promote remodeling of intestinal microbiotas (6, 38–40). Consistent with our findings, *Hungatella* are obesity- and body weight-associated, and enriched in both dietary and surgical weight loss (41–43). The intersection of intestinal microbiotas, antitumor immunity, and cancer outcomes is an area of rapid growth, spurred in part by recent findings that the efficacy of immune checkpoint inhibition can be enhanced by fecal matter transplant (44–46). Indeed, increased representation of *Hungatella* in gut microbiomes following antibiotic treatment has been associated with nonresponse to immune checkpoint inhibition (47).

Prior work from our group and others shows that in obese mice, weight loss induced by VSG, diet switch to a LFD regimen, or CR is associated with reduced mammary tumor growth (5, 33). Here, we delineate the relative efficacy of each intervention in limiting obesity-driven tumor growth. Further, we identify restoration of antitumor immunity markers and reduction of fat mass as common mechanisms potentially underlying the protective effects of weight loss. Our finding that continuous or intermittent CR, had superior efficacy in limiting obesity-driven tumor growth is paralleled in other work showing CR to outperform weight loss driven by ad libitum LFD (5). These findings are particularly germane to the growing interest in the potential for weight loss driven by glucagon-like peptide receptor (GLP1R) and/or glucose-dependent insulinotropic polypeptide receptor (GIPR) agonism to limit obesity-driven tumor growth.

Herein we demonstrate that VSG promotes marked remodeling of epigenetic and transcriptional profiles within adipose tissue, with conserved transcriptional mediators across mouse and human data. We find that VSG, CCR, and ICR each blunt mammary tumor growth in a weight loss- and adiposity loss-dependent manner. Despite the limitation that only one murine model was used, our study comprises comprehensive multi-omic analyses of two independent study cohorts that inform the complex relationships between weight loss, achieved by bariatric surgery or dietary restriction regimens, and the obesity-breast cancer link. Specifically, in a well-established preclinical model of TNBC, we found that i) CCR, ICR, and to a lesser effect VSG, reverse obesity’s procancer effects; ii) shared hallmarks of weight loss-driven reduction of obesity-driven tumor growth in both mice include restoration of transcriptomic signatures of antitumor immunity and marked adipose tissue remodeling; and iii) cecal microbiota changes, particularly enrichment in the genera *Hungatella,* are positively associated with both weight loss and reduced tumor growth.

Given the limitations of bariatric surgery, including risk of adverse effects, high cost, and restrictive third-party insurance coverage (33), further translational exploration of efficacious and sustainable dietary weight loss regimens is urgently needed. Our preclinical findings identify actionable targets and intervention strategies for developing mechanism-based, nonsurgical interventions to reduce the burden of obesity-related breast cancer.

## Methods

### Animals and diets

All animal study protocols were approved and coordinated in compliance with guidelines issued by the University of North Carolina at Chapel Hill Institutional Care and Use Committee (IACUC). Female 8-week-old C57BL/6NCrl mice were purchased from Charles River Laboratories (Wilmington, MA). Upon arrival, mice were group housed on a 12-hour light/dark cycle and offered food and water ad libitum. Following one week of acclimatization, mice were randomized to receive either a low-fat diet (LFD; 10% kcal fat; D12450J, Research Diets) or a high-fat diet (HFD; 60% kcal fat; D12492, Research Diets) to generate normoweight control (CON) or diet-induced obesity (DIO) phenotypes, respectively. Body weights were measured weekly. After 15 weeks on diet, CON mice were continued on LFD (CON-LFD), while DIO mice were randomized to continue the same diet (DIO-HFD) or to undergo weight loss via vertical sleeve gastrectomy (DIO-VSG) or dietary intervention. In Study 1, dietary weight loss was achieved by providing mice ad libitum access to LFD (DIO-LFD). In Study 2, dietary weight loss was achieved by providing mice either a 30% chronic CR diet (DIO-CCR; D15032801 Research Diets), or an intermittent CR diet (DIO-ICR; 14% CR D15032803 and 70% CR D15032804 Research Diets) (5). CCR was accomplished by providing mice 30% fewer daily calories than were consumed by mice in the CON-LFD group.

ICR involved providing mice a 14% CR diet for five days per week and a 70% CR, high-protein diet on two nonconsecutive days per week, thus achieving an average of 30% CR per week relative to the CON-LFD group (5).

### Vertical sleeve gastrectomy and sham procedures

VSG and sham procedures were performed in both Study 1 and Study 2 according to a validated protocol (48). Briefly, VSG involved excision of 70-80% of the lateral stomach. All mice not undergoing VSG underwent a sham procedure to control for physiological insult of surgery. The sham procedure entailed isolation of the stomach and application of gentle, manual pressure using forceps for five seconds. The VSG excision and sham pressure were applied along a line continuous with the esophagus and pylorus. All surgeries occurred within a five-day window. Preoperative fasting, exposure to isoflurane, and administration of analgesics were consistent across all groups. Mice undergoing VSG were postoperatively provided the same LFD as consumed by the control mice to limit the potential for postoperative aversion to high-fat diets and to mimic postoperative diet recommendations for patients undergoing VSG (49).

### Tumor model

Following surgeries, weight loss was closely monitored and once body weights had stabilized (7 weeks for Study 1 and 10 weeks for Study 2), all mice were orthotopically injected with 3.5 x 10^4^ E0771 mammary tumor cells into the fourth mammary fat pad. In vivo tumor growth was monitored by digital calipers. Mice were euthanized by CO2 inhalation four weeks following orthotopic injection. Mammary tumors and tumor-adjacent and contralateral mammary fat pads were weighed, excised, and flash frozen in liquid nitrogen and stored at -80 °C until further analysis. Blood was collected by cardiac puncture, allowed to coagulate for 30 min at room temperature, centrifuged at 1000 rcf for 15 min, and then serum was isolated and stored at -80 °C. To investigate changes in microbiome composition in tumor-naïve mice, fecal samples were collected in Study 1 by applying gentle abdominal pressure immediately prior to VSG surgery and again prior to tumor cell injection and frozen at -20 °C. To investigate changes in microbiome composition of a metabolically important site, cecal samples were collected from Study 2 at euthanasia, flash frozen, and stored at -80 °C.

### Quantitative magnetic resonance imaging (MRI) analysis

Quantitative MRI (Echo Medical Systems, Houston, TX) was used to measure body composition. Lean body mass, fat body mass, and free water were quantified and expressed as percentage of total weight. A randomly selected subset of 8-9 mice per group were analyzed in Study 1 and all mice were analyzed in Study 2.

### Nucleic acid extraction

Total RNA and DNA was extracted from flash frozen tumor and contralateral mammary fat pad samples using TRIzol Reagent (Sigma-Aldrich) according to the manufacturer’s instructions. DNA and RNA integrity was determined by TapeStation analysis (Agilent Technologies).

### Tumor transcriptomic analysis by microarray

Total RNA isolated from tumor tissue was used to synthesize, fragment, and sense-strand label cDNA. cDNA was hybridized to a Mouse Clariom S HT PEG microarray plate (Affymetrix). A GeneTitan MC Instrument (Affymetrix) was used for hybridization, washing, staining, and scanning of the Clariom S peg plate. Data quality control, signal space transformation, and robust multichip average scaling was performed using Transcriptome Analysis Console Software v 4.0.2 (TAC, Thermo Fisher Scientific).

Published transcriptomic profiling of human adipose samples collected either from patients pre- or post-bariatric surgery or gender- and weight-matched controls was accessed through GSE59034 and normalized as above.

### Adipose transcriptomic analysis by RNAseq

Total RNA isolated from murine mammary adipose tissue contralateral to the tumor was used to prepare sequencing libraries with Illumina TruSeq Stranded Total RNA Sample Preparation kit and sequenced using an Illumina HiSeq 2000 instrument (Illumina).

FASTQ files were aligned to the mm10 mouse genome (GRCm38.p4) using STAR v2.4.2 (50) with the following parameters: --outSAMtype BAM Unsorted --quantMode TranscriptomeSAM. Transcript abundance for each sample was estimated with salmon v0.1.19 (51) to quantify the transcriptome defined by Gencode gene annotation. Gene level counts were summed across isoforms, and genes with low expression (defined as fewer than 10 counts across all samples) were removed before downstream analyses. DESeq2 (52) was used to test for differentially expressed genes between interventions.

### Gene set enrichment analysis (GSEA)

GSEA (v4.3.2) was conducted to identify pathways and processes transcriptionally altered by our interventions in both adipose tissue and tumor (29, 30) analysis using SST-RMA normalized microarray data and DESeq2 (52) normalized RNAseq data. Enrichments were calculated for GSEA Hallmarks and Gene Ontology Biological Processes (GOBP). Enrichment mapping was performed to cluster significant (FDRq < 0.05) gene sets by similarity index of genes and to limit redundancy across significant GOBP GSEA results (32).

### DNA methylation analysis by reduced representation bisulfite sequencing (RRBS)

Genome-wide methylation profiles for the mammary fat pad contralateral to the tumor were determined by RRBS. Library preparation and sequencing was performed at the University of North Carolina at Chapel Hill High-Throughput Sequencing Facility.

Alignment and differential methylation analysis was conducted as previously described (53). FASTQ files were aligned to the mm10 mouse genome using Bismark v0.18.1 with default settings (54). BAM files were then sorted with samtools v1.5 (55) and methylation calls were generated using methylKit R package v1.10.0 (56). Bases with low variability (standard deviation of methylation level < 0.05) and extreme read coverage (> 5000X summed across all samples) were removed to avoid PCR artifacts. Data were visualized using nonmetric multidimensional scaling (NMDS) plots. Two outliers (1 CON-LFD, 1 DIO-LFD) were removed from the downstream analysis due to their poor sequencing quality. For RRBS data, methylGSA R package v1.2.3 (57) was used to test for significantly enriched gene sets.

### Adipose tissue oxylipin analysis

Mammary adipose tissue contralateral to the tumor was homogenized and lipids were extracted in methanol, centrifuged at 15,000 rcf for 3 min, and loaded onto an Oasis MAX micro-elution plate (Waters Corp). Oxylipins were washed in methanol and eluted using propanol/acetonitrile (50/50, v/v) containing 5% formic acid. Oxylipins were separated using a Waters Acquity I-Class UPLC and detected using a Waters Xevo TQ-XS triple quadrupole mass spectrometer operating using multiple reaction monitoring in negative ion mode with argon as the collision gas (Waters Corp.). Oxylipins were quantified using the ratio of sample signal to internal isotopically labeled standard peak height and normalized to mass of adipose tissue used.

### 16S rDNA amplicon sequencing

DNA was extracted from fecal and cecal samples following 40 min of vortexing in 0.5 ml of Qiagen PM1 buffer (Qiagen) with 200 mg of 106/500 μm glass beads (Sigma, St. Louis, MO). A KingFisher Flex Purification System was used with Qiagen ClearMag beads to purify DNA (UNC Microbiome Core). 16S 515-806 bp (variable region 4) was amplified from DNA extracted from cecal contents and 16S 27-338 bp (variable region 1-2) was amplified from DNA extracted from fecal samples. PCR amplicons were then sequenced using an Illumina MiSeq platform (Illumina, San Diego, CA) (UNC High-Throughput Sequencing Facility). Sequence variants (SVs) were identified using the DADA2 pipeline (58), with taxonomic classification performed using DADA2 with a SILVA reference database (59). Samples with fewer than 10^4^ total aligned reads and SVs with a frequency <0.01% were removed prior to analysis. Reads were subject to total sum scaling. ⍺ diversity was assessed using Shannon index and observed SVs using MicrobiomeAnalyst (60). β-diversity was assessed using NMDS plots of Bray-Curtis distances calculated using the vegan package (61), and pairwise PERMANOVAs were performed using the pairwiseAdonis package. Association of all genera, assigned using MicrobiomeExplorer (62), with percentage body weight change and tumor mass was performed by Spearman correlation in R, and subject to multiple hypothesis correction.

### Serum secretome analyses

Serum hormone, cytokine, and adipokine concentrations were measured using a Milliplex Mouse Metabolic Hormone Magnetic Bead Panel (MMHMAG-44K), a Bio-Plex Pro^TM^ Mouse Adiponectin Assay, and a 6-Plex Mouse Cytokine Panel (Bio-Rad Laboratories; Hercules, California). Insulin-like growth factor 1 (IGF-1) concentrations were measured using an R&D Systems IGF-1 Bead-Based Single-plex Luminex assay.

### Statistical analysis

One-way analysis of variance (ANOVA) followed by Tukey’s post-hoc test was used to assess the effects of diet and weight loss across groups. For ɑ-diversity measures Kruskal-Wallis or Mann-Whitney U tests were used. PERMANOVA was used to assess average Bray-Curtis distances between groups. Correction for multiple testing was achieved using the Benjamini-Hochberg procedure for all transcriptomic, epigenetic, and 16S rDNA amplicon sequencing. Results were analyzed using GraphPad Prism software (GraphPad Software Inc.) and R version 3.4.3. *P* ≤ 0.05 was considered statistically significant.

### Material availability

Microarray, RNA-Seq, and RRBS datasets have been deposited in the NCBI Gene Expression Omnibus under the accession number GSE230474. All other data are available from authors on reasonable request.

## Author Contributions

Conceptualization: MFC, KKC, ELR, SDH. Methodology: MFC, KKC, ELR, AGL, FF, WG, JSP, RJS, AAF. Software: MFC, KKC, TLM, SSD, FF, WG, JSP, AAF. Validation:

MFC, KKC. Formal analysis: MFC, KKC, TLM, SSD, FF, WG, JSP, AAF. Investigation: MFC, KKC, ELR, TLM, SSD, ETR, AL, SAK, AJP, LAS, LWB, EMG, GLM. Data

Curation: MFC, KKC, WG, EMG. Writing - Original Draft: MFC, KKC, EMG, SDH. Writing - Review & Editing: MFC, KKC, LWB, FF, EMG, JSP, IMC, RJS, SDH. Visualization: MFC, SSD. Supervision: JSP, RJS, AAF, SDH. Funding acquisition: SDH.

## Acknowledgements

This work was supported by a grant from the National Institutes of Health (R35 CA197627) and the Breast Cancer Research Foundation (BCRF-21-073) to SDH. The authors would like to acknowledge Dr. Emily L Rossi for the important contributions she made in establishing the bariatric surgery protocol used in the reported studies.

## References

1. WCRF/AICR. Diet, nutrition, physical activity and cancer: a global perspective. Continuous update project expert report. 2018.

2. Devericks EN, Carson MS, McCullough LE, Coleman MF, and Hursting SD. The obesity-breast cancer link: a multidisciplinary perspective. Cancer and Metastasis Reviews. 2022;41(3):607–25.

3. Hales CM, Carroll MD, Fryar CD, and Ogden CL. Prevalence of Obesity and Severe Obesity Among Adults: United States, 2017-2018. NCHS Data Brief. 2020(360):1–8.

4. Bohm MS, Sipe LM, Pye ME, Davis MJ, Pierre JF, and Makowski L. The role of obesity and bariatric surgery-induced weight loss in breast cancer. Cancer and Metastasis Reviews. 2022;41(3):673–95.

5. Bowers LW, Doerstling SS, Shamsunder MG, Lineberger CG, Rossi EL, Montgomery SA, et al. Reversing the Genomic, Epigenetic, and Triple-Negative Breast Cancer–Enhancing Effects of Obesity. Cancer Prevention Research. 2022;15(9):581–94.

6. Bowers LW, Glenny EM, Punjala A, Lanman NA, Goldbaum A, Himbert C, et al. Weight Loss and/or Sulindac Mitigate Obesity-associated Transcriptome, Microbiome, and Protumor Effects in a Murine Model of Colon Cancer. Cancer Prev Res (Phila). 2022;15(8):481–95.

7. Aminian A, Wilson R, Al-Kurd A, Tu C, Milinovich A, Kroh M, et al. Association of Bariatric Surgery With Cancer Risk and Mortality in Adults With Obesity. JAMA. 2022;327(24):2423–33.

8. Lunger F, Aeschbacher P, Nett PC, and Peros G. The impact of bariatric and metabolic surgery on cancer development. Front Surg. 2022;9:918272.

9. Sjöström L, Gummesson A, Sjöström CD, Narbro K, Peltonen M, Wedel H, et al. Effects of bariatric surgery on cancer incidence in obese patients in Sweden (Swedish Obese Subjects Study): a prospective, controlled intervention trial. The Lancet Oncology. 2009;10(7):653–62.

10. Tsui ST, Yang J, Zhang X, Docimo S, Jr., Spaniolas K, Talamini MA, et al. Development of cancer after bariatric surgery. Surg Obes Relat Dis.2020;16(10):1586–95.

11. Alalwan AA, Friedman J, Park H, Segal R, Brumback BA, and Hartzema AG. US national trends in bariatric surgery: A decade of study. Surgery. 2021;170(1):13–7.

12. Welbourn R, Hollyman M, Kinsman R, Dixon J, Liem R, Ottosson J, et al. Bariatric Surgery Worldwide: Baseline Demographic Description and One-Year Outcomes from the Fourth IFSO Global Registry Report 2018. Obesity Surgery. 2019;29(3):782–95.

13. Cummings DE, and Rubino F. Metabolic surgery for the treatment of type 2 diabetes in obese individuals. Diabetologia. 2018;61(2):257–64.

14. Cummings DE, Arterburn DE, Westbrook EO, Kuzma JN, Stewart SD, Chan CP, et al. Gastric bypass surgery vs intensive lifestyle and medical intervention for type 2 diabetes: the CROSSROADS randomised controlled trial. Diabetologia. 2016;59(5):945–53.

15. Ospanov O, Akilzhanova A, Buchwald JN, Fursov A, Bekmurzinova F, Rakhimova S, et al. Stapleless vs Stapled Gastric Bypass vs Hypocaloric Diet: a Three-Arm Randomized Controlled Trial of Body Mass Evolution with Secondary Outcomes for Telomere Length and Metabolic Syndrome Changes. Obes Surg. 2021;31(7):3165–76.

16. Ribaric G, Buchwald JN, and McGlennon TW. Diabetes and weight in comparative studies of bariatric surgery vs conventional medical therapy: a systematic review and meta-analysis. Obes Surg. 2014;24(3):437–55.

17. Freedhoff Y, and Hall KD. Weight loss diet studies: we need help not hype. The Lancet. 2016;388(10047):849–51.

18. Anderson AS, Renehan AG, Saxton JM, Bell J, Cade J, Cross AJ, et al. Cancer prevention through weight control—where are we in 2020? British Journal of Cancer. 2021;124(6):1049–56.

19. Iyengar NM, Gucalp A, Dannenberg AJ, and Hudis CA. Obesity and Cancer Mechanisms: Tumor Microenvironment and Inflammation. Journal of Clinical Oncology. 2016;34(35):4270–6.

20. Scheithauer TPM, Rampanelli E, Nieuwdorp M, Vallance BA, Verchere CB, van Raalte DH, et al. Gut Microbiota as a Trigger for Metabolic Inflammation in Obesity and Type 2 Diabetes. Frontiers in Immunology. 2020;11.

21. Ahmed T, Das SK, Golden JK, Saltzman E, Roberts SB, and Meydani SN. Calorie restriction enhances T-cell-mediated immune response in adult overweight men and women. J Gerontol A Biol Sci Med Sci. 2009;64(11):1107–13.

22. Dyck L, Prendeville H, Raverdeau M, Wilk MM, Loftus RM, Douglas A, et al. Suppressive effects of the obese tumor microenvironment on CD8 T cell infiltration and effector function. J Exp Med. 2022;219(3).

23. Gibson JT, Orlandella RM, Turbitt WJ, Behring M, Manne U, Sorge RE, et al. Obesity-Associated Myeloid-Derived Suppressor Cells Promote Apoptosis of Tumor-Infiltrating CD8 T Cells and Immunotherapy Resistance in Breast Cancer. Frontiers in immunology. 2020;11:590794-.

24. Wang Z, Aguilar EG, Luna JI, Dunai C, Khuat LT, Le CT, et al. Paradoxical effects of obesity on T cell function during tumor progression and PD-1 checkpoint blockade. Nat Med. 2019;25(1):141–51.

25. Naaman SC, Shen S, Zeytinoglu M, and Iyengar NM. Obesity and Breast Cancer Risk: The Oncogenic Implications of Metabolic Dysregulation. The Journal of Clinical Endocrinology & Metabolism. 2022;107(8):2154–66.

26. Delahanty LM, Wadden TA, Goodwin PJ, Alfano CM, Thomson CA, Irwin ML, et al. The Breast Cancer Weight Loss trial (Alliance A011401): A description and evidence for the lifestyle intervention. Obesity. 2022;30(1):28–38.

27. Willems AEM, Sura-de Jong M, van Beek AP, Nederhof E, and van Dijk G. Effects of macronutrient intake in obesity: a meta-analysis of low-carbohydrate and low-fat diets on markers of the metabolic syndrome. Nutr Rev. 2021;79(4):429–44.

28. Harvie M, and Howell A. Behavioral Sciences. 2017.

29. Subramanian A, Tamayo P, Mootha VK, Mukherjee S, Ebert BL, Gillette MA, et al. Gene set enrichment analysis: a knowledge-based approach for interpreting genome-wide expression profiles. Proc Natl Acad Sci U S A. 2005;102(43):15545-50.

30. Liberzon A, Birger C, Thorvaldsdottir H, Ghandi M, Mesirov JP, and Tamayo P. The Molecular Signatures Database (MSigDB) hallmark gene set collection. Cell Syst. 2015;1(6):417–25.

31. The Gene Ontology resource: enriching a GOld mine. Nucleic Acids Res. 2021;49(D1):D325–d34.

32. Reimand J, Isserlin R, Voisin V, Kucera M, Tannus-Lopes C, Rostamianfar A, et al. Pathway enrichment analysis and visualization of omics data using g:Profiler, GSEA, Cytoscape and EnrichmentMap. Nat Protoc. 2019;14(2):482–517.

33. Sipe LM, Chaib M, Korba EB, Jo H, Lovely MC, Counts BR, et al. Response to immune checkpoint blockade improved in pre-clinical model of breast cancer after bariatric surgery. Elife. 2022;11.

34. Kondo K, Ishigaki Y, Gao J, Yamada T, Imai J, Sawada S, et al. Bach1 deficiency protects pancreatic β-cells from oxidative stress injury. Am J Physiol Endocrinol Metab. 2013;305(5):E641–8.

35. Dehondt H, Marino A, Butruille L, Mogilenko DA, Nzoussi Loubota AC, Chávez- Talavera O, et al. Adipocyte-specific FXR-deficiency protects adipose tissue from oxidative stress and insulin resistance and improves glucose homeostasis. Molecular Metabolism. 2023:101686.

36. Catrysse L, and van Loo G. Inflammation and the Metabolic Syndrome: The Tissue-Specific Functions of NF-κB. Trends in Cell Biology. 2017;27(6):417–29.

37. Bhardwaj P, and Brown KA. Obese Adipose Tissue as a Driver of Breast Cancer Growth and Development: Update and Emerging Evidence. Frontiers in Oncology. 2021;11.

38. Debédat J, Clément K, and Aron-Wisnewsky J. Gut Microbiota Dysbiosis in Human Obesity: Impact of Bariatric Surgery. Curr Obes Rep. 2019;8(3):229–42.

39. Ilhan ZE, DiBaise JK, Isern NG, Hoyt DW, Marcus AK, Kang DW, et al. Distinctive microbiomes and metabolites linked with weight loss after gastric bypass, but not gastric banding. Isme j. 2017;11(9):2047–58.

40. Fouladi F, Brooks AE, Fodor AA, Carroll IM, Bulik-Sullivan EC, Tsilimigras MCB, et al. The Role of the Gut Microbiota in Sustained Weight Loss Following Roux-en-Y Gastric Bypass Surgery. Obes Surg. 2019;29(4):1259–67.

41. Aron-Wisnewsky J, Prifti E, Belda E, Ichou F, Kayser BD, Dao MC, et al. Major microbiota dysbiosis in severe obesity: fate after bariatric surgery. Gut. 2019;68(1):70.

42. Sbierski-Kind J, Grenkowitz S, Schlickeiser S, Sandforth A, Friedrich M, Kunkel D, et al. Effects of caloric restriction on the gut microbiome are linked with immune senescence. Microbiome. 2022;10(1):57.

43. Dang JT, Mocanu V, Park H, Laffin M, Hotte N, Karmali S, et al. Roux-en-Y gastric bypass and sleeve gastrectomy induce substantial and persistent changes in microbial communities and metabolic pathways. Gut Microbes. 2022;14(1):2050636.

44. Baruch EN, Youngster I, Ben-Betzalel G, Ortenberg R, Lahat A, Katz L, et al. Fecal microbiota transplant promotes response in immunotherapy-refractory melanoma patients. Science. 2021;371(6529):602–9.

45. Gopalakrishnan V, Spencer CN, Nezi L, Reuben A, Andrews MC, Karpinets TV, et al. Gut microbiome modulates response to anti-PD-1 immunotherapy in melanoma patients. Science. 2018;359(6371):97–103.

46. Vetizou M, Pitt JM, Daillere R, Lepage P, Waldschmitt N, Flament C, et al. Anticancer immunotherapy by CTLA-4 blockade relies on the gut microbiota. Science. 2015;350(6264):1079–84.

47. Hakozaki T, Richard C, Elkrief A, Hosomi Y, Benlaïfaoui M, Mimpen I, et al. The Gut Microbiome Associates with Immune Checkpoint Inhibition Outcomes in Patients with Advanced Non-Small Cell Lung Cancer. Cancer Immunol Res. 2020;8(10):1243–50.

48. Wilson-Perez HE, Chambers AP, Ryan KK, Li B, Sandoval DA, Stoffers D, et al. Vertical sleeve gastrectomy is effective in two genetic mouse models of glucagon-like Peptide 1 receptor deficiency. Diabetes. 2013;62(7):2380–5.

49. Ratner C, Shin JH, Dwibedi C, Tremaroli V, Bjerregaard A, Hartmann B, et al. Anorexia and Fat Aversion Induced by Vertical Sleeve Gastrectomy Is Attenuated in Neurotensin Receptor 1-Deficient Mice. Endocrinology. 2021;162(9):bqab130.

50. Dobin A, Davis CA, Schlesinger F, Drenkow J, Zaleski C, Jha S, et al. STAR: ultrafast universal RNA-seq aligner. Bioinformatics. 2013;29(1):15–21.

51. Patro R, Duggal G, Love MI, Irizarry RA, and Kingsford C. Salmon provides fast and bias-aware quantification of transcript expression. Nature Methods. 2017;14(4):417–9.

52. Love MI, Huber W, and Anders S. Moderated estimation of fold change and dispersion for RNA-seq data with DESeq2. Genome Biol. 2014;15(12):550.

53. Rossi EL, de Angel RE, Bowers LW, Khatib SA, Smith LA, Van Buren E, et al. Obesity-Associated Alterations in Inflammation, Epigenetics, and Mammary Tumor Growth Persist in Formerly Obese Mice. Cancer Prev Res (Phila). 2016;9(5):339–48.

54. Krueger F, and Andrews SR. Bismark: a flexible aligner and methylation caller for Bisulfite-Seq applications. Bioinformatics. 2011;27(11):1571–2.

55. Li H, Handsaker B, Wysoker A, Fennell T, Ruan J, Homer N, et al. The Sequence Alignment/Map format and SAMtools. Bioinformatics. 2009;25(16):2078–9.

56. Akalin A, Kormaksson M, Li S, Garrett-Bakelman FE, Figueroa ME, Melnick A, et al. methylKit: a comprehensive R package for the analysis of genome-wide DNA methylation profiles. Genome Biology. 2012;13(10):R87.

57. Ren X, and Kuan PF. methylGSA: a Bioconductor package and Shiny app for DNA methylation data length bias adjustment in gene set testing. Bioinformatics. 2019;35(11):1958–9.

58. Callahan BJ, McMurdie PJ, Rosen MJ, Han AW, Johnson AJA, and Holmes SP. DADA2: High-resolution sample inference from Illumina amplicon data. Nature Methods. 2016;13(7):581–3.

59. Quast C, Pruesse E, Yilmaz P, Gerken J, Schweer T, Yarza P, et al. The SILVA ribosomal RNA gene database project: improved data processing and web- based tools. Nucleic Acids Res. 2013;41(Database issue):D590-6.

60. Chong J, Liu P, Zhou G, and Xia J. Using MicrobiomeAnalyst for comprehensive statistical, functional, and meta-analysis of microbiome data. Nature Protocols. 2020;15(3):799–821.

61. Oksanen J SG, Blanchet F, Kindt R, Legendre P, Minchin P, O’Hara R, Solymos P, Stevens M, Szoecs E,, Wagner H BM, Bedward M, Bolker B, Borcard D, Carvalho G, Chirico M, De Caceres M, Durand S,, Evangelista H FR, Friendly M, Furneaux B, Hannigan G, Hill M, Lahti L, McGlinn D, Ouellette M,, and Ribeiro Cunha E ST, Stier A, Ter Braak C, Weedon J 2022.

62. Reeder J, Huang M, Kaminker JS, and Paulson JN. MicrobiomeExplorer: an R package for the analysis and visualization of microbial communities. Bioinformatics. 2021;37(9):1317–8.

