## Supplemental Figures for "Calorie Restriction Outperforms Bariatric Surgery in a Murine Model of Obesity and Triple-Negative Breast Cancer"

### Fig. S1

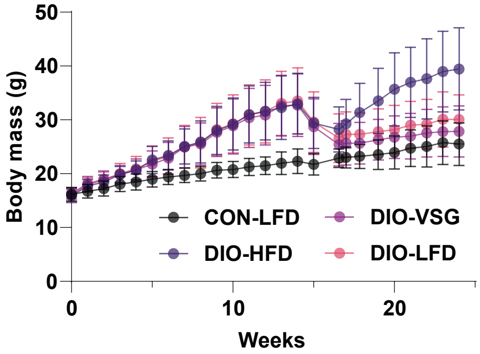

Body mass (g)
50
40
30
20
10
0
0
10
20
Weeks
CON-LFD
DIO-HFD
DIO-VSG
DIO-LFD

### Fig. S2

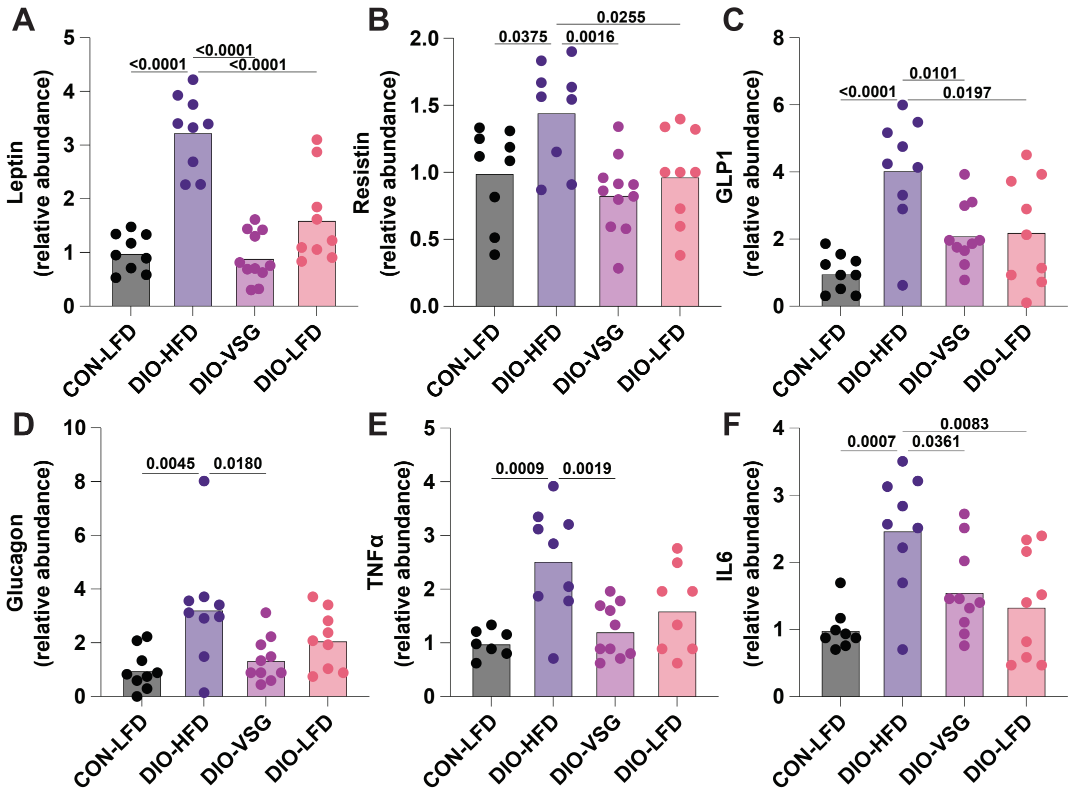

A
B
C
D
E
F
Leptin (relative abundance)
Resistin (relative abundance)
GLP1 (relative abundance)
Glucagon (relative abundance)
TNFα (relative abundance)
IL6 (relative abundance)
<0.0001
<0.0001
<0.0001
0.0375
0.0016
0.0255
0.0101
0.0197
0.0045
0.0180
0.0009
0.0019
0.0007
0.0361
0.0083
CON-LFD
DIO-HFD
DIO-VSG
DIO-LFD

### Fig. S3

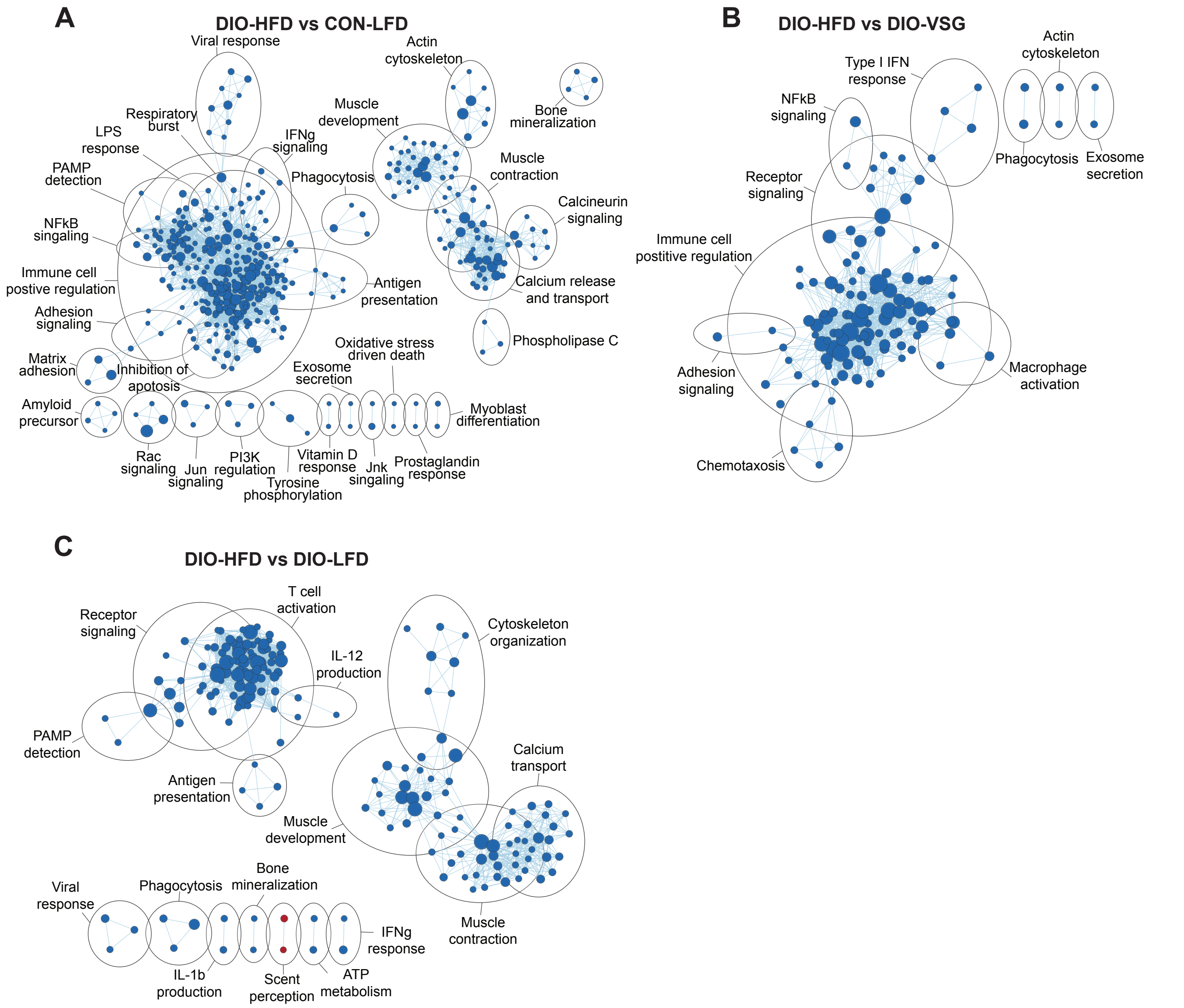
A
DIO-HFD vs CON-LFD
Viral response
Actin cytoskeleton
Bone mineralization
Respiratory burst
LPS response
IFNg signaling
Muscle development
Muscle contraction
PAMP detection
Phagocytosis
Calcineurin signaling
NFkB singaling
Immune cell postive regulation
Antigen presentation
Calcium release and transport
Adhesion signaling
Phospholipase C
Matrix adhesion
Inhibition of apotosis
Oxidative stress driven death
Exosome secretion
Amyloid precursor
Myoblast differentiation
Rac signaling
Jun signaling
PI3K regulation
Vitamin D response
Jnk singaling
Prostaglandin response
Tyrosine phosphorylation
B
DIO-HFD vs DIO-VSG
Actin cytoskeleton
Type I IFN response
NFkB signaling
Phagocytosis
Exosome secretion
Receptor signaling
Immune cell postitive regulation
Adhesion signaling
Macrophage activation
Chemotaxosis
C
DIO-HFD vs DIO-LFD
T cell activation
Receptor signaling
Cytoskeleton organization
IL-12 production
PAMP detection
Calcium transport
Antigen presentation
Muscle development
Bone mineralization
Viral response
Phagocytosis
IFNg response
Muscle contraction
IL-1b production
Scent perception
ATP metabolism

### Fig. S4

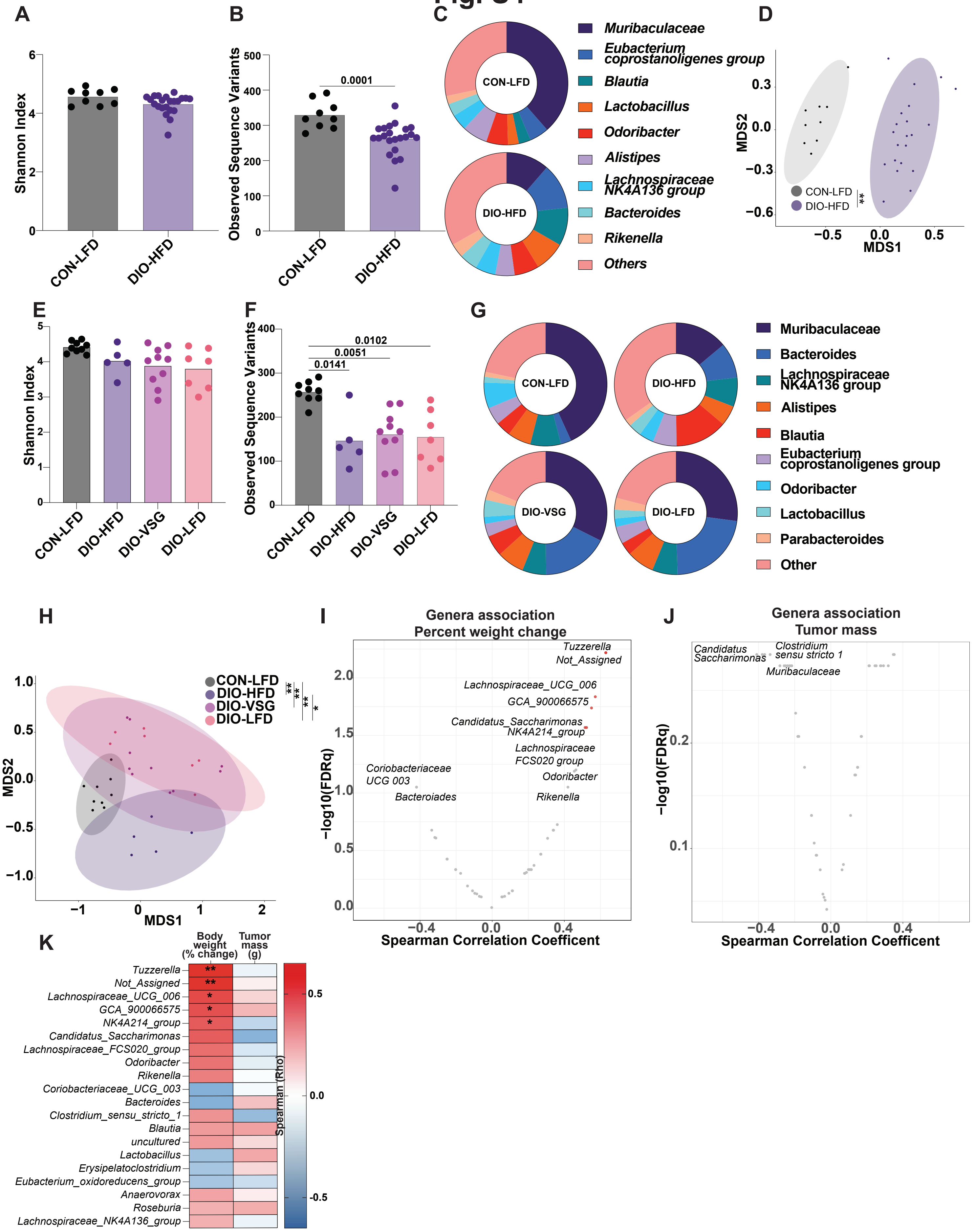
A
Shannon Index
CON-LFD
DIO-HFD
B
Observed Sequence Variants
0.0001
CON-LFD
DIO-HFD
C
CON-LFD
DIO-HFD
Muribaculaceae
Eubacterium coprostanoligenes group
Blautia
Lactobacillus
Odoribacter
Alistipes
Lachnospiraceae NK4A136 group
Bacteroides
Rikenella
Others
D
MDS2
MDS1
CON-LFD
DIO-HFD
**
E
Shannon Index
CON-LFD
DIO-HFD
DIO-VSG
DIO-LFD
F
Observed Sequence Variants
0.0102
0.0051
0.0141
G
CON-LFD
DIO-HFD
DIO-VSG
DIO-LFD
Muribaculaceae
Bacteroides
Lachnospiraceae NK4A136 group
Alistipes
Blautia
Eubacterium coprostanoligenes group
Odoribacter
Lactobacillus
Parabacteroides
Other
H
CON-LFD
DIO-HFD
DIO-VSG
DIO-LFD
MDS2
MDS1
I
Genera association
Percent weight change
Tuzzerella
Not_Assigned
Lachnospiraceae_UCG_006
GCA_900066575
Candidatus_Saccharimonas
NK4A214_group
Lachnospiraceae FCS020 group
Odoribacter
Rikenella
Coriobacteriaceae UCG 003
Bacteroiades
−log10(FDRq)
Spearman Correlation Coefficent
J
Genera association
Tumor mass
Candidatus Saccharimonas
Clostridium sensu stricto 1
Muribaculaceae
−log10(FDRq)
Spearman Correlation Coefficent
K
Body weight (% change)
Tumor mass (g)
Tuzzerella
Not_Assigned
Lachnospiraceae_UCG_006
GCA_900066575
NK4A214_group
Candidatus_Saccharimonas
Lachnospiraceae_FCS020_group
Odoribacter
Rikenella
Coriobacteriaceae_UCG_003
Bacteroides
Clostridium_sensu_stricto_1
Blautia
uncultured
Lactobacillus
Erysipelatoclostridium
Eubacterium_oxidoreducens_group
Anaerovorax
Roseburia
Lachnospiraceae_NK4A136_group
Spearman (Rho)

### Fig. S5

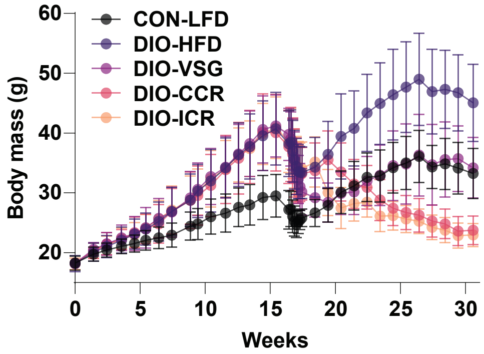

CON-LFD
DIO-HFD
DIO-VSG
DIO-CCR
DIO-ICR
Body mass (g)
60
50
40
30
20
0
5
10
15
20
25
30
Weeks

**Figure S.1**

Body mass of all animals over course of experiment n=21 CON-LFD, 21 DIO-HFD, 24 DIO-VSG, 19 DIO-LFD.

**Figure S.2**

(A-F) Circulating adipokines determined by multiplex ELISA. n=9 CON-LFD, 9 DIO-HFD, 11 DIO-VSG, 9 DIO-LFD. One-way ANOVA with Tukey's post hoc test.

**Figure S.3**

Enrichment maps of significant (FDRq<0.05) GOBP gene sets for pairwise comparisons with DIO-HFD. (A) DIO-HFD vs. CON-LFD, (B) DIO-HFD vs. DIO-VSG, and (C) DIO-HFD vs. DIO-LFD. Node size reflects gene set size, line weight reflects overlap coefficient (minimum 0.5), blue color denotes enriched in comparison relative to DIO-HFD.

**Figure S.4**

(A) Observed sequence variants and (B) Shannon index of pre-intervention fecal microbial communities. (C) Relative contribution of the 10 most frequent genera to each group pre-intervention. (D) NMDS plot of Bray-Curtis distances of pre-intervention microbial communities. (E) Observed sequence variants and (F) Shannon diversity of post-intervention fecal microbial communities. (G) Relative contribution of the 10 most frequent genera to each group post-intervention. (H) NMDS plot of Bray-Curtis distances of post-intervention microbial communities. Spearman correlation between all genera and (I) percent body weight and (J) tumor mass. (K) Spearman correlation coefficients of the 20 genera showing the highest correlation coefficients with percent body weight change and tumor mass.

(A-D) n=9 CON-LFD and 21 DIO-HFD. (E-K) n=9 CON-LFD, 5 DIO-HFD, 10 DIO-VSG, and 7 DIO-LFD.

**Figure S.5**

Body mass of all animals over course of experiment n=20 CON-LFD, 18 DIO-HFD, 14 DIO-VSG, 19 DIO-ICR, 16 DIO-CCR.
